# Iron Status Influences Mitochondrial Disease Progression in Complex I-Deficient Mice

**DOI:** 10.1101/2021.09.29.462431

**Authors:** Anthony S. Grillo, CJ Kelly, Vivian T. Ha, Camille M. Bodart, Sydney Huff, Reid K. Couch, Nicole T. Herrel, Hyunsung D. Kim, Azaad O. Zimmermann, Jessica Shattuck, Yu-Chen Pan, Matt Kaeberlein

**Affiliations:** Department of Laboratory Medicine & Pathology, University of Washington, Seattle, United States

## Abstract

Mitochondrial dysfunction caused by aberrant Complex I assembly and reduced activity of the electron transport chain is pathogenic in many genetic and age-related diseases. Mice missing the Complex I subunit NADH dehydrogenase [ubiquinone] iron-sulfur protein 4 (NDUFS4) are a leading mammalian model of severe mitochondrial disease that exhibit many characteristic symptoms of Leigh Syndrome including oxidative stress, neuroinflammation, brain lesions, and premature death. NDUFS4 knockout mice have decreased expression of nearly every Complex I subunit. As Complex I normally contains at least 8 iron-sulfur clusters and more than 25 iron atoms, we asked whether a deficiency of Complex I may lead to intracellular iron perturbations thereby accelerating disease progression. Consistent with this, iron supplementation accelerates symptoms of brain degeneration in these mice while iron restriction delays the onset of these symptoms and increases survival. NDUFS4 knockout mice display signs of iron overload in the liver including increased expression of hepcidin, and show changes in iron responsive element-regulated proteins consistent with increased intracellular iron that were prevented by iron restriction. These results suggest that perturbed iron homeostasis may contribute to pathology in Leigh Syndrome and possibly other mitochondrial disorders.

## Introduction

Inherited mitochondrial defects cause several lethal mitochondriopathies such as Leigh Syndrome, MELAS, and Friedreich’s Ataxia (*1*). Reduced assembly and/or activity of respiratory Complex I or other complexes of the electron transport chain (ETC) are widely implicated in the etiology of most of these mitochondrial diseases (*1-4*). Additionally, a common feature of age-related diseases, including Alzheimer’s disease, Parkinson’s disease, heart disease, and diabetes, is decreased oxidative phosphorylation through aberrant ETC function (*5-9*). Complex I is the largest of the ETC complexes and is made up of 45 subunit proteins that represent a considerable portion of the protein mass of the inner mitochondrial membrane (*10*). It coordinates the transfer of electrons from NADH to ubiquinone via a shuttle of 8 or more redox-active iron-sulfur clusters found on the peripheral arm in the mitochondrial matrix, but little is known on the effects of improper regulation, assembly, biogenesis, and dynamics of these iron-sulfur clusters in the pathophysiology of Complex I deficiencies.

One of the most prevalent hereditary mitochondrial diseases is Leigh Syndrome, which is characterized by lactic acidosis, neuroinflammation, brain lesions of the basal ganglia, and death within the first few years of life (*11*). Nearly 35% of Leigh Syndrome cases can be caused by various mutations affecting Complex I including NDUFS4 and several other iron sulfur proteins on the redox-active peripheral arm (*1, 12*). Mice missing the Complex I subunit NDUFS4 are a leading mammalian model of Leigh Syndrome (*13*). We previously reported knockout of the iron-sulfur protein NDUFS4 (*Ndufs4*^*-/-*^) in mice decreases expression of nearly all Complex I subunits, and we were unable to observe appreciable Complex I or respiratory supercomplex formation (*14, 15*). Iron-sulfur cluster deficiencies in cells have recently been linked to an accumulation of iron (*16*), and prior evidence in cells suggests inhibition of Complex I with rotenone promotes iron accumulation that is dependent on iron sensor protein activity (*17-20*). However, altered iron metabolism and its improper regulation because of deficiencies of an iron-sulfur cluster protein in *Ndufs4*^*-/-*^ mice is unknown.

Iron as an essential nutrient is the most biologically abundant transition metal that plays key roles in physiology (*21-23*). It is pervasively utilized as an enzymatic co-factor due to its unique readiness to undergo facile redox cycling in cellular milieu. Iron is largely localized to mitochondria due to its essential role in electron transfer during cellular respiration and in mitochondrial metabolism (*24*). The high reactivity of iron, however, facilitates reactive oxygen species generation, oxidative stress, ferroptosis, and organ damage when in excess (*25-27*). Iron homeostasis is tightly controlled through multiple transcriptional, translational, and post-translational iron-dependent feedback regulatory mechanisms such as the iron-responsive element (IRE) signaling pathway and the hepcidin-ferroportin axis (*21, 22*). However, abnormal iron accumulation and/or utilization can overwhelm these regulatory mechanisms leading to disease (*22*). Here we utilized the *Ndufs4*^*-/-*^ mice to test the hypothesis that Complex I deficiencies may alter normal cellular or regional iron distribution which contributes to mitochondrial disease progression.

## Results

### Iron Status Underlies Disease Progression in NDUFS4-KO Mice

To evaluate the influence of iron on disease progression in *Ndufs4*^*-/-*^ mice, we first observed the onset of clasping in mice treated with the FDA-approved iron chelator deferiprone. Clasping is a common feature in the early stages of brain degeneration in these mice that immediately precedes severe neuroinflammation, weight loss, and ataxia (*15*). There is high correlation between the onset of clasping and observed lifespan (Fig. 1A), making this neurobehavioral symptom an appropriate readout of disease progression. Control *Ndufs4*^*-/-*^ mice fed standard chow began clasping around 40 days of age, consistent with our prior reports (*14*). To probe the role of iron in neurodegeneration, we added the FDA-approved iron chelator deferiprone to the water of *Ndufs4*^*-/-*^ mice after weaning. We observed treatment with this brain-penetrating iron chelator delayed the onset of clasping (Fig. 1B and Fig. Supp. 1A). Deferiprone treatment also increased median lifespan in *Ndufs4*^*-/-*^ mice (Fig. 1C and Fig. Supp. 1B, C).

**Figure 1.**
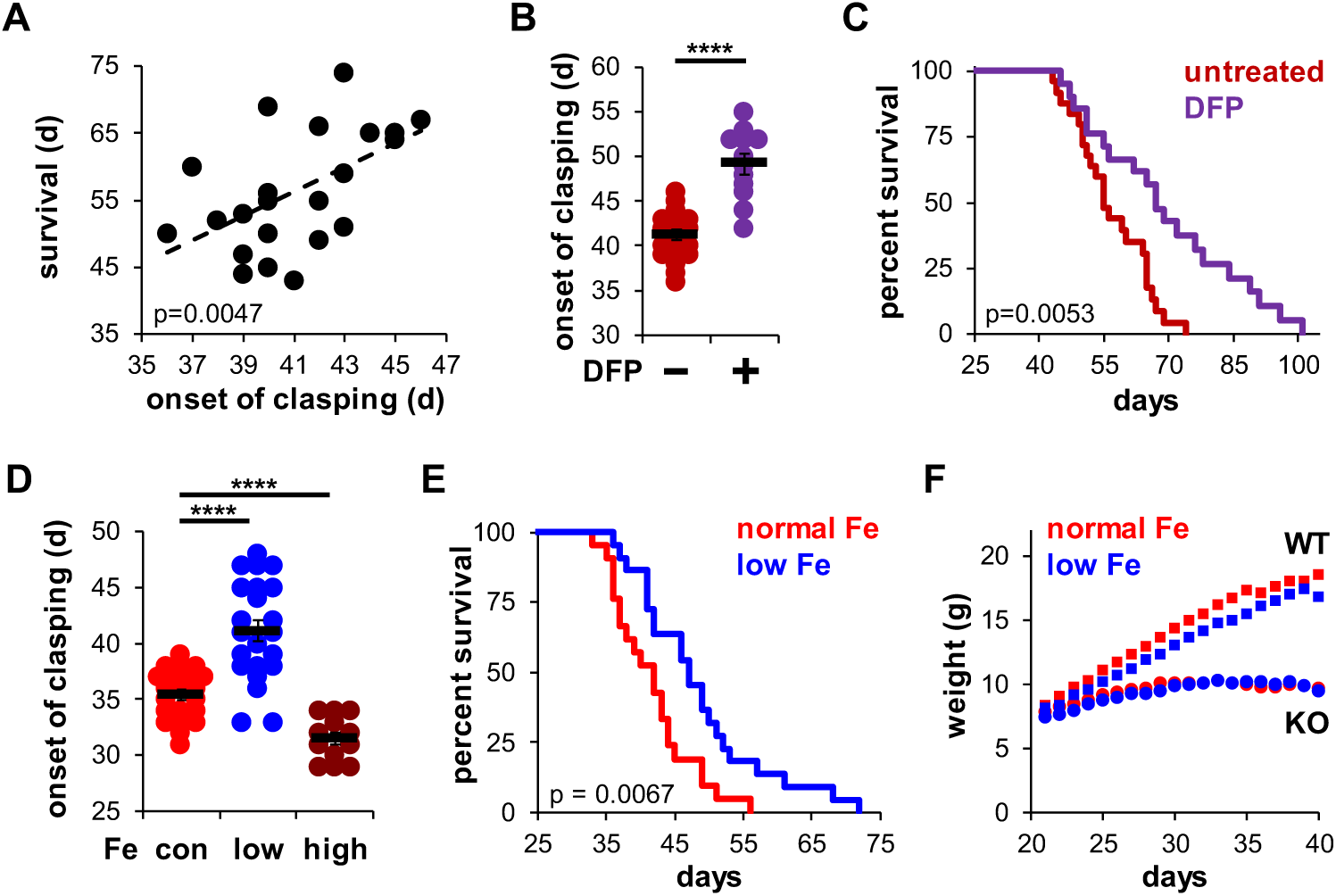
(A) Correlation between the onset of clasping and survival. Each point represents data from a single mouse. p=0.0047, Pearson’s test. (B) Age at which *Ndufs4*^*-/-*^ mice exhibited the clasping phenotype on chow diet. Mice were treated with either vehicle or deferiprone (DFP) in the water (2 mg/mL) from weaning. (C) Survival curves of *Ndufs4*^*-/-*^ mice fed a chow diet and treated with deferiprone in the water (2 mg/mL) from weaning. (D) Onset of clasping in *Ndufs4*^*-/-*^ mice on AIN-93G synthetic diet containing normal (40 ppm, con) or low (8 ppm) iron starting from weaning. Mice on control diet (40 ppm Fe) were also treated with iron-dextran (100 mg/kg every 3 days via i.p. injection, high) from weaning. (E) Survival curves of mice on normal (40 ppm) or low (8 ppm) AIN-93G synthetic diet. (F) Weight gain in wild type (WT, square markers) or *Ndufs4*^*-/-*^ mice (KO, circle markers) on AIN-93G synthetic diet containing normal (40 ppm, red) or low (8 ppm, blue) concentrations of iron. p-value was calculated by log-rank for lifespan analyses. **** p<0.0001, t-test.

Because robust quantification of iron clearance is difficult to achieve due to an unreliable variability of iron in non-synthetic chows, we performed similar experiments using a synthetic AIN-93G control diet containing normal levels of iron (40 ppm). It was reported that *Ndufs4*^*-/-*^ mice fed a synthetic diet clasp at earlier ages and have shorter lifespans than *Ndufs4*^*-/-*^ mice fed standard chow diets (*28*), which we similarly observed (Fig. 1D, E). High iron supplementation via intraperitoneal injection of iron-dextran (100 mg/kg every 3 days) accelerated the onset of clasping (Fig. 1D), while feeding mice a low iron (8 ppm) synthetic diet dramatically delayed the onset of clasping without causing any significant deleterious changes in weight (Fig. 1D and 1F). This delay in disease progression was associated with ∼15% increase in lifespan (Fig. 1E).

We next performed hematological analysis of blood isolated from these mice. As expected, *Ndufs4*^*-/-*^ mice fed a low iron diet showed signs of microcytic, hypochromic anemia consistent with iron deficiency. We observed decreased hematocrit, reduced hemoglobin levels, and decreased red blood cell production in *Ndufs4*^*-/-*^ mice fed a low iron diet compared to the normal iron diet cohorts (Fig. 2A-C). We also observed decreases in MCV, MCH, and MCHC with the low iron diet (Fig. 2D-F).

**Figure 2.**
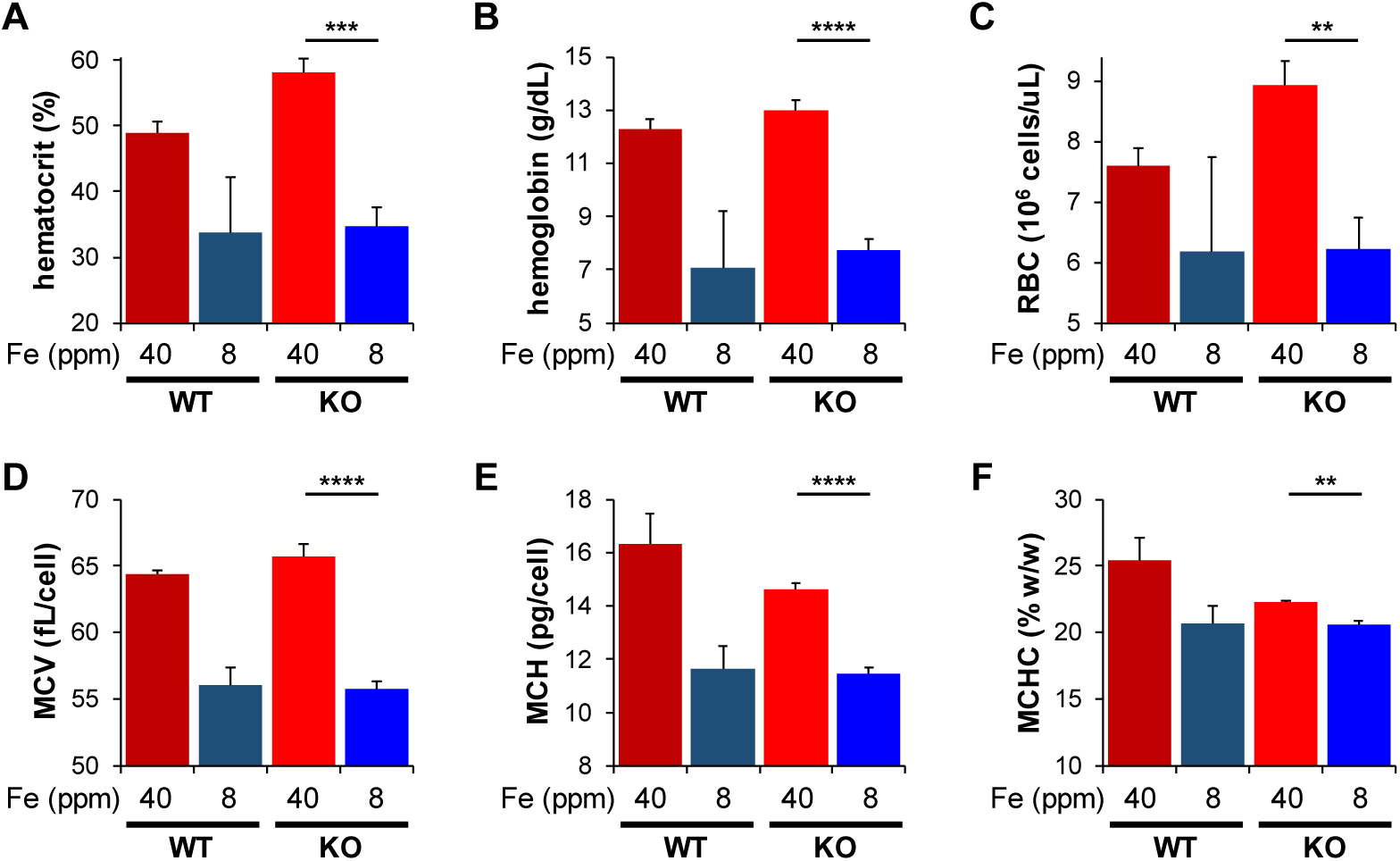
Complete blood count in PND30 WT or *Ndufs4*^*-/-*^ mice on normal (40 ppm) or low (8 ppm) AIN-93G synthetic diet to quantify (A) hematocrit, (B) hemoglobin, (C) red blood cell count which were used to calculate (D) mean corpuscular volume, (E) mean corpuscular hemoglobin, and (F) mean corpuscular hemoglobin concentration. N=3-6 mice. ** p<0.01, *** p<0.001, **** p<0.0001, t-test.

### Metal Imbalances in Tissues of NDUFS4-KO Mice

We collected liver and other tissues from WT and *Ndufs4*^*-/-*^ mice near the onset of clasping (i.e. PND35) to quantify total iron levels by inductively coupled plasma mass spectrometry (ICP-MS). This time point was chosen as it is near the median age that control *Ndufs4*^*-/-*^ mice begin clasping (Fig. 1D). ICP-MS of tissue digestates revealed that *Ndufs4*^*-/-*^ mice fed the control synthetic diet had ∼3-fold increases in total liver iron relative to WT mice (Fig. 3A-G and Table 1). Significantly decreased iron levels in kidney and duodenum were observed in mice fed the low iron diet (Fig. 3A-G and Table 1), and we observed a global reduction in iron levels in the tissues tested (Fig. 4A). Iron restriction did not affect the relative weights of these tissues compared to mice fed the control diet (Fig. Supp. 2). We observed no differences in total iron in whole brain digestates from 35-day old WT and *Ndufs4*^*-/-*^ mice fed the normal or low iron diets (Fig. 3B and Table 1).

**Figure 3.**
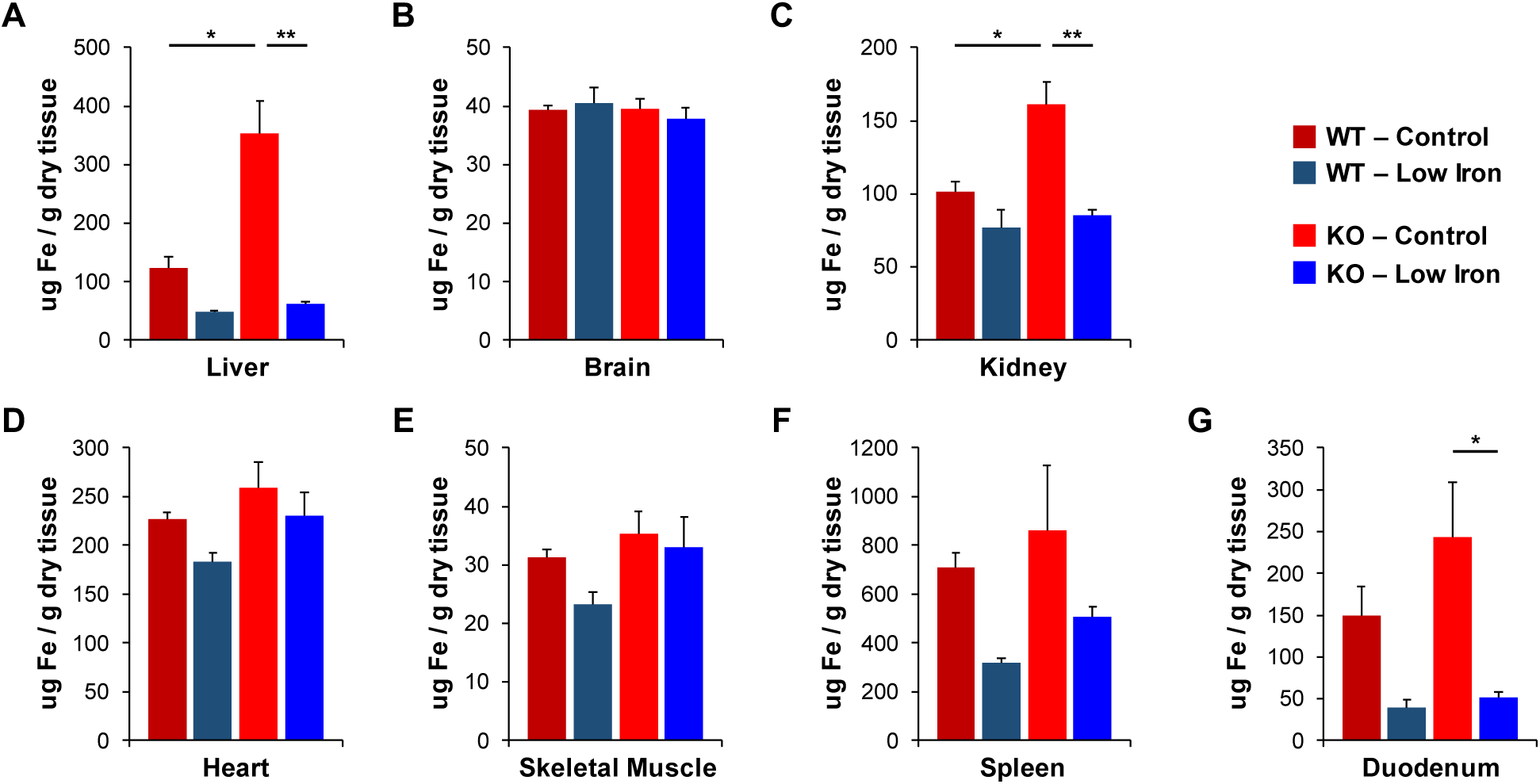
Quantification of total iron by ICP-MS from WT and *Ndufs4*^*-/-*^ mice at PND35 fed control (40 ppm) or low (8 ppm) AIN-93G in (A) liver, (B) whole brain, (C) kidney, (D) heart, (E) quadricep, (F) spleen, and (G) duodenum. N=4-6 mice. * p<0.05, ** p<0.01, t-test.

**Table 1.** ICP-MS quantification of biologically relevant metals in WT and *Ndufs4*^*-/-*^ tissues at PND35 fed a normal (40 ppm) and low (8 ppm) iron AIN-93G synthetic diet. Metals were measured as µg metal relative to total dry weight of tissue. N=4-6 mice, - p<0.10, * p<0.05, ** p<0.01, *** p<0.001, t-test.

|  | Metal<br>(µg/g) | *Ndufs4*<sup>+/+</sup> Mice |  | *Ndufs4*<sup>-/-</sup> Mice |  | p value |  |
| --- | --- | --- | --- | --- | --- | --- | --- |
|  |  | AIN-93G<br>Normal Fe | AIN-93G<br>Low Iron | AIN-93G<br>Normal Iron | AIN-93G<br>Low Iron | WT-Con<br>vs KO-Con | KO-Con<br>vs KO-Low |
| Liver | Fe | 122.9 ± 18.9 | 48.0 ± 3.5 | 352.5 ± 55.5 | 62.5 ± 3.6 | * | ** |
|  | Mn | 2.20 ± 0.33 | 5.68 ± 0.63 | 3.70 ± 0.44 | 8.62 ± 0.51 | * | *** |
|  | Zn | 47.5 ± 5.07 | 62.0 ± 5.8 | 60.9 ± 5.0 | 71.2 ± 4.0 | - |  |
|  | Cu | 9.6 ± 0.64 | 12.7 ± 0.9 | 13.9 ± 1.5 | 15.6 ± 1.2 | * |  |
| Brain | Fe | 39.3 ± 0.78 | 40.6 ± 2.5 | 39.6 ± 1.8 | 37.8 ± 1.9 |  |  |
|  | Mn | 1.46 ± 0.04 | 1.94 ± 0.05 | 1.71 ± 0.09 | 1.91 ± 0.09 | - |  |
|  | Zn | 37.7 ± 0.53 | 48.3 ± 1.6 | 44.1 ± 1.8 | 49.1 ± 3.1 | * |  |
|  | Cu | 10.3 ± 0.52 | 11.4 ± 0.5 | 11.8 ± 0.4 | 12.1 ± 0.7 | - |  |
| Kidney | Fe | 101.9 ± 6.4 | 77.1 ± 12.4 | 161.5 ± 15.4 | 85.8 ± 3.5 | * | ** |
|  | Mn | 4.54 ± 0.44 | 5.53 ± 0.31 | 4.48 ± 0.20 | 6.01 ± 0.51 |  | - |
|  | Zn | 55.8 ± 1.8 | 56.6 ± 2.6 | 58.9 ± 4.8 | 58.4 ± 5.7 |  |  |
|  | Cu | 15.1 ± 0.3 | 15.3 ± 0.9 | 17.5 ± 0.7 | 18.0 ± 0.9 | * |  |
| Heart | Fe | 226.8 ± 6.9 | 182.7 ± 9.5 | 259.2 ± 26.9 | 230.7 ± 24.0 |  |  |
|  | Mn | 2.51 ± 0.09 | 3.00 ± 0.05 | 2.59 ± 0.19 | 3.72 ± 0.35 |  | ** |
|  | Zn | 46.6 ± 3.0 | 47.7 ± 1.5 | 31.5 ± 3.0 | 41.3 ± 5.3 | * |  |
|  | Cu | 30.5 ± 1.2 | 30.5 ± 0.1 | 31.7 ± 2.9 | 40.4 ± 2.8 |  | - |
| Skeletal<br>Muscle | Fe | 31.2 ± 1.4 | 23.3 ± 2.1 | 35.4 ± 3.9 | 33.0 ± 5.3 |  |  |
|  | Mn | 0.53 ± 0.05 | 0.83 ± 0.08 | 0.65 ± 0.08 | 0.88 ± 0.10 |  |  |
|  | Zn | 20.7 ± 0.7 | 21.8 ± 1.4 | 22.5 ± 2.8 | 18.8 ± 1.8 |  |  |
|  | Cu | 3.80 ± 0.24 | 3.85 ± 0.22 | 4.60 ± 0.56 | 4.40 ± 0.44 |  |  |
| Spleen | Fe | 710.4 ± 59.1 | 316.9 ± 22.3 | 860.5 ± 265.6 | 507.1 ± 43.7 |  |  |
|  | Mn | 0.94 ± 0.09 | 1.28 ± 0.08 | 0.92 ± 0.40 | 1.45 ± 0.23 |  |  |
|  | Zn | 110.5 ± 7.5 | 123.1 ± 10.4 | 109.0 ± 38.7 | 126.0 ± 6.8 |  |  |
|  | Cu | 5.13 ± 0.55 | 6.43 ± 0.43 | 3.84 ± 1.26 | 5.29 ± 0.57 |  |  |
| Duodenum | Fe | 150.1 ± 34.2 | 40.0 ± 8.6 | 243.0 ± 66.3 | 51.7 ± 6.2 |  | * |
|  | Mn | 6.78 ± 1.2 | 12.88 ± 2.63 | 6.76 ± 1.28 | 17.1 ± 1.7 |  | ** |
|  | Zn | 89.2 ± 2.2 | 84.9 ± 12.7 | 101.4 ± 9.6 | 114.1 ± 9.3 |  |  |
|  | Cu | 9.4 ± 0.3 | 8.4 ± 1.3 | 10.8 ± 0.9 | 11.2 ± 0.8 |  |  |

**Figure 4.**
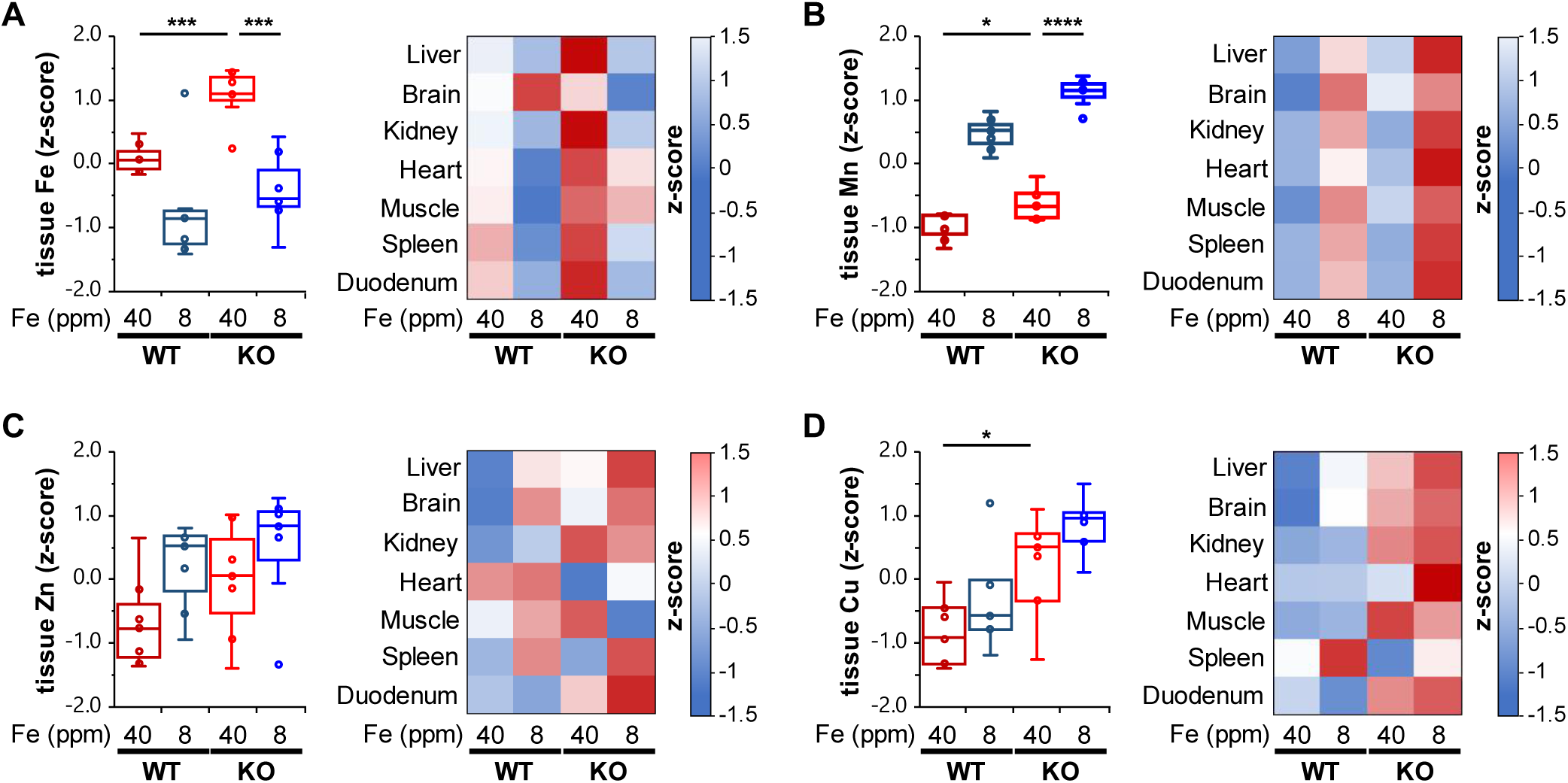
(A) Left, Box and whisker plot of combined z-score-normalized ICP-MS values of tissue iron, (B) manganese, (C) zinc, and (D) copper in WT and *Ndufs4*^*-/-*^ mice on control (40 ppm) or low (8 ppm) AIN-93 diet (each point represents z-score value for an individual tissue). (A-D) Right, heat map of individualized z-scores by tissue. N=4-6 mice. * p<0.05, *** p<0.001, **** p<0.0001, t-test.

Similarities between divalent metals, notably iron (II) and manganese (II), lead to low stoichiometric selectivity of divalent metal transport proteins for these metals (*29*). These transport proteins often have higher affinities for copper or manganese. The biological selectivity, however, is primarily controlled by the relative availability of these metals in the cell (*23, 30-32*). Labile iron is most often in ∼10-fold or greater concentrations relative to manganese, copper, or other metals in biological systems, and thus iron is often preferred. In cases of iron restriction, however, the observed selectivity for transport through metal transport proteins shifts to prefer manganese. Manganese metabolism is sensitive to changes in iron homeostasis because of the promiscuity of many iron transport proteins for other divalent metals (*29*).

We thus quantified total levels of manganese and other metals in WT and *Ndufs4*^*-/-*^ mice on normal or low iron diet by ICP-MS. Consistent with the putative effect iron restriction has on increasing manganese uptake, we observed elevated manganese levels in WT and *Ndufs4*^*-/-*^ mice fed a low iron diet in the liver, brain, kidney, and other tissues relative to control mice (Fig. 4B and Table 1). The control *Ndufs4*^*-/-*^ mice had increased basal manganese levels compared to WT cohorts. This may be due to a compensatory upregulation of mitochondrial manganese-dependent superoxide dismutase (MnSOD aka SOD2) to combat iron-mediated oxidative stress when mice were fed the normal iron diet (*33*). We did not observe any notable changes in copper, zinc, or other biorelevant metals in iron restricted mice (Table 1 and Fig. 4C, D). However, *Ndufs4*^*-/-*^ mice fed the normal iron diet showed increased copper levels relative to control WT mice (Fig. 4D).

### Increased Oxidative Stress in NDUFS4-KO Mice

Excess free iron often accelerates the accumulation of reactive oxygen species by reacting with molecular oxygen through the Fenton reaction. *Ndufs4*^*-/-*^ mice have been reported to have elevated, and presumably toxic, levels of oxygen in the brain, blood, and other tissues due to their decreased respiratory activity (*34*). The unmitigated accumulation of reactive oxygen species causes oxidative stress and damages cells through the radical-mediated peroxidation of polyunsaturated fatty acids (PUFAs). These lipid peroxyl radicals break down into malondialdehyde (MDA) and 4-hydroxynonenal and promote local inflammation.

We first asked whether *Ndufs4*^*-/-*^ mice on control iron diet had increased non-heme iron relative to WT controls using a ferrozine assay. Ferrozine forms a purple-colored chelate with iron allowing for the colorimetric detection and quantification of free or weakly bound (e.g. non-heme) iron. This free iron can react with O_2_ and PUFAs to generate ROS. Consistent with this, we observed livers from *Ndufs4*^*-/-*^ mice on the control iron diet had increased non-heme iron (Fig. 5A).

**Figure 5.**
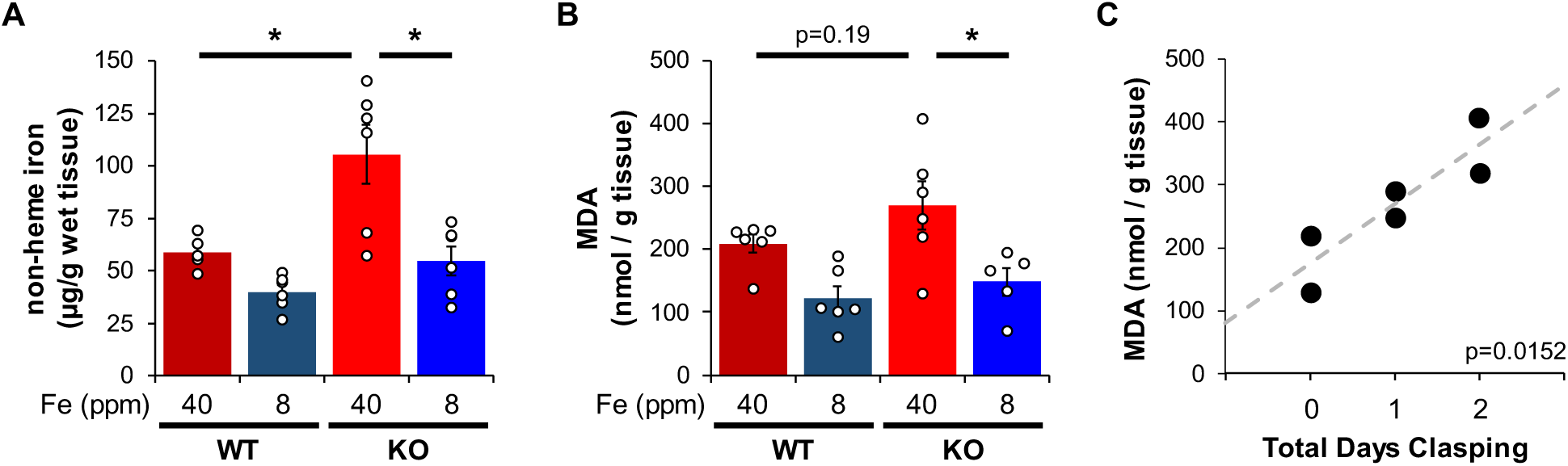
(A) Quantification of non-heme iron by ferrozine assay and (B) MDA-TBA adduct in livers from WT and *Ndufs4*^*-/-*^ mice at PND35 that were fed control (40 ppm) or low (8 ppm) AIN-93G synthetic diet. (C) Correlation between days since *Ndufs4*^*-/-*^ mice began displaying the clasping phenotype with detected liver MDA levels from (B). p=0.0152, Pearson’s test. N=5-6 mice. * p<0.05, t-test.

This led us to next ask whether livers from *Ndufs4*^*-/-*^ mice had increased PUFA oxidation using MDA as a readout. We quantified MDA levels using a TBARS assay, in which thiobarbituric acid reacts with MDA to form a TBA-MDA adduct that can be quantified colorimetrically. We observed a trend toward higher MDA levels in livers from control *Ndufs4*^*-/-*^ mice compared to WT cohorts (Fig. 5B), with a strong correlation between the length of time *Ndufs4*^*-/-*^ mice exhibited neurodegenerative symptoms (i.e. clasping) and observed MDA levels (Fig. 5C). Iron-deficient mice showed decreased MDA levels (Fig. 5B). Collectively, this data supports the hypothesis that iron accumulation in livers of *Ndufs4*^*-/-*^ mice promotes oxidative stress.

### Iron Responsive Element Iron Regulation Suggests Increased Labile Iron

Increased MDA production suggests increased intracellular labile iron, but it is challenging to directly probe changes in intracellular iron distribution *in vivo*. However, the expression of iron-dependent regulatory proteins can be used as an effective readout. Body iron distribution is solely controlled by regulating iron absorption, metabolism, and storage as there are no known adaptive biological mechanisms for iron excretion (*27*). The expression of iron regulatory proteins is highly sensitive to intracellular iron perturbations. For example, upregulation of the iron storage protein ferritin helps reduce cellular labile iron and avoid oxidative stress in cases of iron overload. Regulation is primarily achieved at the translational and post-translational levels through iron-responsive elements (IREs) and the hepcidin-ferroportin axis (*21, 22*).

IREs are found in the 5’ or 3’-untranslated regions of mRNA coding for proteins involved in iron uptake (e.g. transferrin receptor 1), iron storage (e.g. ferritin), and cellular iron export (e.g. ferroportin). The iron regulatory proteins (IRPs) bind to these IREs and either block translation (5’-IRE) or prevent endonuclease-mediated mRNA degradation (3’-IRE) upstream of the poly-A tail (Fig. Supp. 3). Recognition of labile iron by IRPs induces a conformational change that prevents IRPs binding to the IREs. Holo-IRP1 adopts aconitase activity in the cytosol while IRP2 is ubiquitinated and degraded with iron overload.

To probe changes in intracellular iron status, we evaluated mRNA levels and expression of proteins known to be involved in IRE-dependent regulation. Expression of the cytosolic iron storage protein ferritin is controlled by 5’-IREs found on its 24 heavy (*Fth1*) and light (*Ftl1*) chain subunits and is an excellent readout of free cellular iron. Ferritin can deposit more than 4,000 iron atoms into its central core, thereby acting as an iron sponge (*35*). In cases of iron overload, IRPs dissociate from the 5’-IRE allowing for translation of FTH1 and FTL1 (Fig. Supp. 3). Consistent with iron overload in livers of control *Ndufs4*^*-/-*^ mice, we observed knockout of NDUFS4 increased FTH1 protein expression (Fig. 6A and 6B). Iron restriction in *Ndufs4*^*-/-*^ mice downregulated FTH1 expression, consistent with iron deficiency anemia (Fig. 6A and 6B).

**Figure 6.**
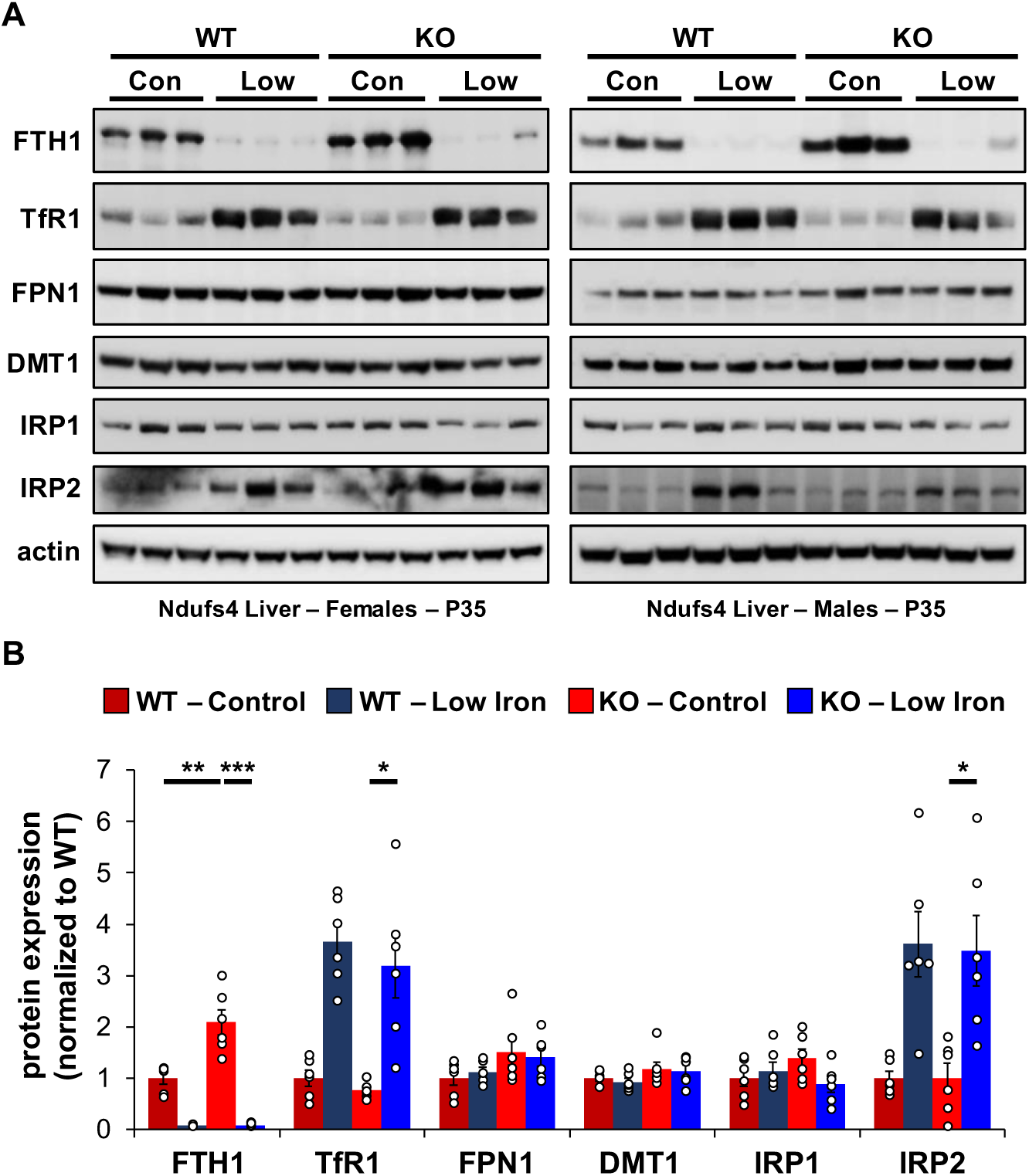
(A) Representative western blot images and (B) densitometry (relative to actin) of proteins involved in regulation of iron transport, storage, or metabolism in livers from PND35 WT and *Ndufs4*^*-/-*^ mice fed a control (40 ppm) or low (8 ppm) iron AIN-93G synthetic diet from weaning. Each lane represents protein extract from a single mouse. * p<0.05, ** p<0.01, *** p<0.001, t-test.

Transferrin Receptor 1 (TFR1) mediates the cellular internalization of iron-bound transferrin. *Tfr1* mRNA contains a 3’-IRE and is normally expressed at high levels with iron restriction. To adapt to iron overload, IRPs dissociate from the *Tfr1* mRNA leading to its endonuclease-mediated degradation (Fig. Supp. 3). Control *Ndufs4*^*- /-*^ mice showed decreased levels of *Tfr1* mRNA in liver by qRT-PCR relative to WT mice (Fig. 7A), consistent with increased labile iron in the mitochondrial disease mice. As expected, we observed upregulation of *Tfr1* mRNA and TFR1 protein in livers from the WT and *Ndufs4*^*-/-*^ mice fed the low iron diet (Fig. 6A, 6B, and 7A). We also observed changes in other IRE-regulated genes at the mRNA or protein level in liver as expected for IRE-dependent regulation (Fig. 6A, 6B, and 7A-E). Iron deficiency in *Ndufs4*^*-/-*^ mice prevented the altered expression of these genes (Fig. 7A-E), consistent with reduced intracellular iron. We did not observe appreciable differences in FTH1 or TFR1 expression in the brain (Fig. Supp. 4).

**Figure 7.**
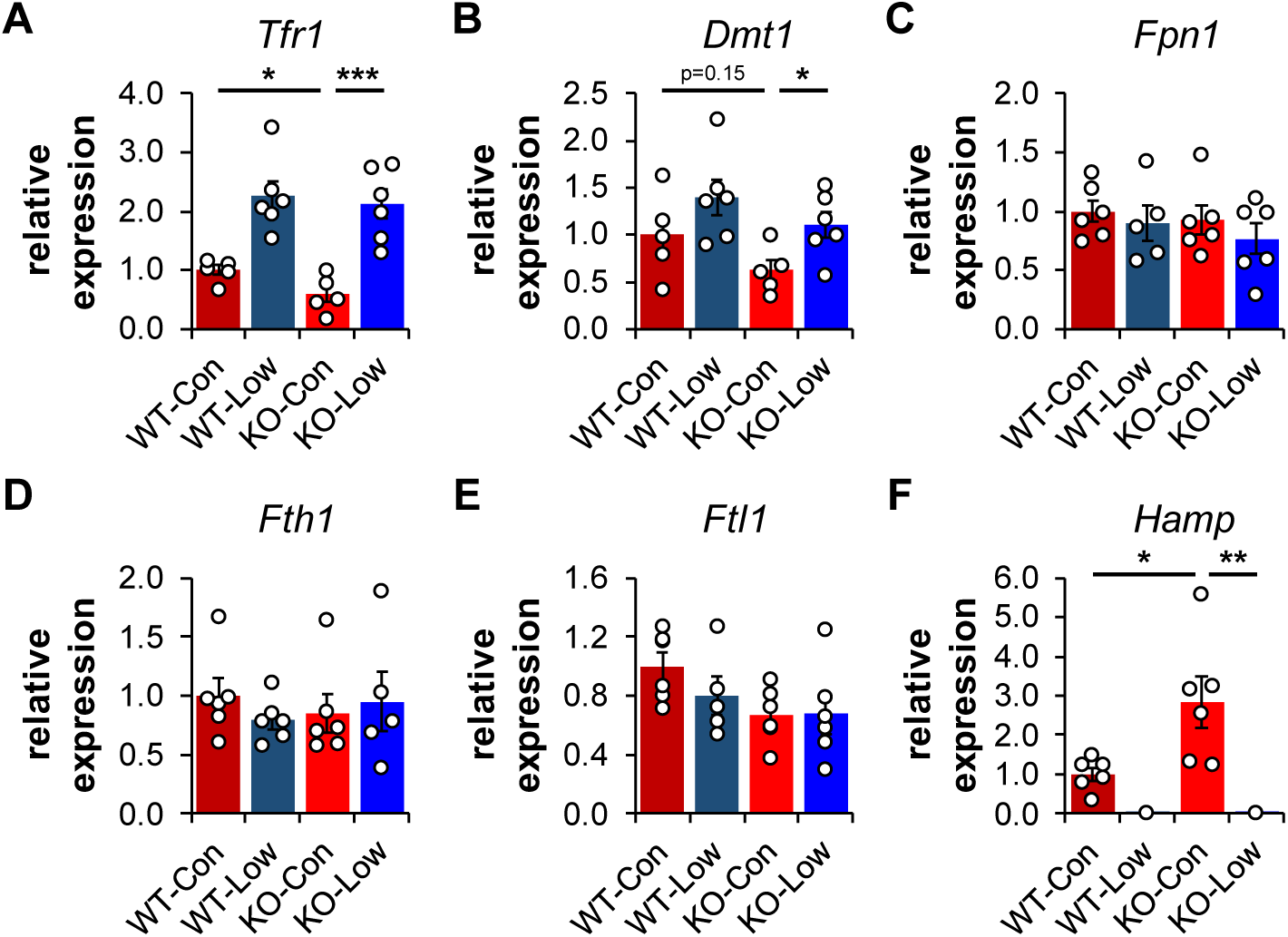
(A) Quantification of relative mRNA expression of *Tfr1*, (B) *Dmt1*, (C) *Fpn1*, (D) *Fth1* (ferritin heavy chain 1), (E) *Ftl1* (ferritin light chain 1), and (F) *Hamp* (hepcidin) in livers from PND35 WT and *Ndufs4*^*-/-*^ (KO) mice fed a control (40 ppm) and low (8 ppm) iron AIN-93G synthetic diet from weaning. * p<0.05, ** p<0.01, *** p<0.001, t-test.

### The Hepcidin-Ferroportin Axis in a Complex I Deficiency

The peptide hepcidin is a liver-derived master regulator of iron absorption, iron recycling, iron storage, and in erythropoiesis (*21*). In situations of excess iron, hepatic production and systemic circulation of hepcidin promotes ubiquitination and proteasomal degradation of the iron export protein ferroportin (FPN1) thereby trapping iron in duodenal epithelia and in liver macrophages. This promotes the storage of excess iron in the liver and reduces iron absorption from the diet as an effective adaptive mechanism protecting against iron-mediated damage in cases of iron overload. While *Fpn1* mRNA contains a 5’-IRE, hepcidin-mediated degradation of FPN1 prevents its IRE-dependent upregulation in the liver or gut epithelia with iron overload. Consistent with this, we did not observe significant increases in FPN1 protein in *Ndufs4*^*-/-*^ liver (Fig. 6A and 5B).

We thus asked whether hepcidin production increased in livers of *Ndufs4*^*-/-*^ mice. We quantified *Hamp1* mRNA encoding the hepcidin peptide (Fig. 7F). Livers of control *Ndufs4*^*-/-*^ mice had 3-fold increased transcript levels of *Hamp1*, further consistent with our ferritin data suggesting increased hepatic iron stores. Iron restriction drastically downregulated *Hamp1* transcription in liver to nearly undetectable levels (Fig. 7F).

## Discussion

### Pathogenic Iron Dyshomeostasis in Diverse Age-Related and Genetic Diseases

Iron is widely recognized as a critical mediator in the progression of many age-related and genetic neurodegenerative diseases (*27*). There is increased interest in recent years to better understand its pathogenic role in neurodegeneration with brain iron accumulation and other age-related disorders (*36*). For example, iron deposition in the basal ganglia of Parkinson’s Disease (PD) patients and mouse models accelerates the aggregation of alpha-synuclein, promotes free radical accumulation, increases lipid peroxidation, and is implicated in dementia and motor impairment (*36*). Treatment with the Complex I inhibitor rotenone is widely used as a PD model that reproduces many features including oxidative damage, dopaminergic degeneration in the substantia nigra, and Lewy Body inclusions (*18*). Rotenone-treatment increases the mitochondrial and cytosolic labile iron pool in neurons *in vitro* and in animal models (*17-20*).

Several mitochondrial disorders caused by deficiencies of other ETC complexes or assembly factors can result in clinical symptoms overlapping with Leigh Syndrome. For example, point mutations of the Complex III chaperone ubiquinol-cytochrome c reductase complex chaperone (BCS1L) can cause the rare hereditary disease GRACILE (<u>G</u>rowth <u>R</u>etardation, <u>A</u>minoaciduria, <u>C</u>holestasis, Iron overload, <u>L</u>actic acidosis, and <u>E</u>arly death) syndrome (*37-41*). BCS1L normally mediates the transfer of Rieske iron-sulfur protein into the pre-assembled Complex III dimer, but its deficiency leads to elevated liver iron, increased serum iron or ferritin, and hypotransferrinemia (*37*). There is considerable overlap between the metabolic alterations in patients with Leigh Syndrome and GRACILE syndrome. This is demonstrated by findings that some mutations of BCS1L cause Leigh Syndrome (e.g. P99L) (*42*) while other mutations in this protein alternatively cause GRACILE syndrome (e.g. S78G) (*40*). Similar metabolic changes include decreased respiratory supercomplex formation, reduced O_2_ consumption, high amino acid levels, NAD^+^ dyshomeostasis, increased glycolysis, and increased lactate fermentation (*37-41*). As seen in Leigh Syndrome, GRACILE Syndrome patients with encephalopathy experience seizures, developmental delays, and other neurodegenerative symptoms (*42, 43*). These shared phenotypic manifestations of Leigh and GRACILE Syndrome may be a result of overlapping pathophysiology of iron accumulation.

### Iron Accumulation in Various Tissues

Our ICP-MS data quantifying metals in several tissues strongly suggests that there is an accumulation of iron in the *Ndufs4*^*-/-*^ mice (Fig. 3 and Table 1). Interestingly, we see the most drastic changes in the liver. The liver plays a central role in iron absorption, metabolism, and regulation by acting as the primary iron storage vessel to protect sensitive organs, such as the brain. The liver promotes the controlled release of iron into the blood to maintain non-toxic physiological levels, is largely responsible for synthesizing circulating iron-binding proteins such as transferrin, and produces hepcidin – the master regulating peptide of iron absorption – in response to changes in hepatic iron stores (*21, 27*). Because of the liver’s critical importance in maintaining systemic iron, our results suggest there is an excess of body iron in the setting of this Complex I deficiency. While the *Ndufs4*^*-/-*^ mice experience a severe neurometabolic disease, we previously reported that liver-specific knockout of S6K1, but not neuronal S6K1, can delay the onset of disease and prolong survival (*44*). Our data further implicates aberrations in liver physiology in *Ndufs4*^*-/-*^ mouse disease progression that may translate to patients with Leigh Syndrome.

We were unable to detect any noticeable differences in iron in whole brain digestates by ICP-MS (Fig. 3). The reason for this is unclear, however, it may be due to well-controlled brain-specific adaptive mechanisms that limit iron absorption through the blood-brain barrier (*45*), or due to differential iron distribution in regions that are unaffected in *Ndufs4*^*-/-*^ mice (*46*). For example, mice missing the iron regulatory protein IRP2 show neurocognitive deficits consistent with brain iron overload, but iron is only deposited in glial cells and white matter while neuronal iron contrastingly decreases (*47, 48*). Age-progressive iron accumulation is primarily found in the basal ganglia and other regions that control motor function while whole brain iron deposition is largely unchanged (*49*). In support of this, *Ndufs4*^*-/-*^ pathology is commonly observed in the basal ganglia, cerebellum, and olfactory bulbs. These brain regions tend to have higher iron levels than other regions (*36, 49*). Iron does not need to cross the blood-brain-barrier to accumulate in the olfactory bulb or brain stem, and as such these regions are more sensitive to disturbances in iron homeostasis. We observed that dietary iron restriction, and treatment with an iron chelator, both delay the onset of neurological symptoms (i.e. clasping). This is consistent with the idea that altered whole body iron status can influence neurodegeneration.

### Cellular Iron Distribution Perturbations as a Result of a Complex I Deficiency

Our data does not elucidate the specific forms or oxidation state of cellular iron that accumulates in *Ndufs4*^*-/-*^ mice, but it is possible that the speciation of iron is altered. As Complex I normally utilizes >25 iron atoms in its iron-sulfur clusters, its deficiency could alter the iron composition in the cell. In support of this, recent research shows iron-sulfur cluster deficiencies cause iron imbalances (*16*). In contrast to protein-bound iron, excess cellular labile iron is widely associated with ROS generation, lipid peroxidation, oxidative stress, ferroptosis, or other damaging processes (*25*). We observed increased iron (Fig. 5A), increased levels of the lipid peroxidation byproduct MDA (Fig. 5B), and changes in the IRE-associated proteins TFR1 and ferritin subunits (Fig. 6A). This data collectively supports increased free cytosolic iron in *Ndufs4*^*-/-*^ mouse liver that is recognized by IRP1 and IRP2.

Iron is especially localized to the mitochondria due to its widespread use in energy production, metabolism, and normal ROS signaling. A large proportion of this is normally utilized in oxidative phosphorylation. It is thus interesting to consider whether the distribution of iron is perturbed not only at the organ level, but also intracellularly, in *Ndufs4*^*-/-*^ mice. A recent report in rotenone-treated SH-SY5Y neuroblastoma cells showed Complex I inhibition increased both cytosolic and mitochondrial labile iron by calcein-green and RPA fluorescence, respectively (*18*). As NDUFS4 is found in mitochondria, alterations in subcellular iron distribution remains an attractive avenue of pursuit in the setting of a Complex I deficiency.

### The effect of iron deficiency on oxygen status

Recent reports provide some evidence that *Ndufs4*^*-/-*^ mice are hyperoxic, show increased PO_2_ levels in the brain and blood, and have perturbed redox status (*34*). This excess of oxygen is implicated in disease etiology. Consistent with this, housing mice in low oxygen atmospheres (11% O_2_) corrects abnormal gas tensions, decreases mitochondrial H_2_O_2_ production, prevents brain lesions, and considerably extends lifespan (*50, 51*). Induction of severe anemia by regular early-life phlebotomy every other day in combination with an iron deficient diet is similarly beneficial (*34*). However, the content of this iron-restricted diet, the characterization of iron status, and the influence of iron on disease progression in *Ndufs4*^*-/-*^ mice were unclear. Our data does not indicate whether the low iron diet decreases oxygen tension to delay the onset of neurodegenerative symptoms, but it may play a role. However, the extent of lifespan extension with a low iron diet is small in comparison to severe anemia by combining iron restriction with regular phlebotomy. Both may be necessary to drastically reduce oxygen delivery to the brain.

It was previously reported that genetic activation of the hypoxia-inducible factor (HIF) regulatory pathway via knockout of prolyl hydroxylases, nestin, or Von Hippel-Lindau protein in *Ndufs4*^*-/-*^ mice is detrimental (*34*). Jain and Zazzeron *et al*. reported median lifespan in control *Ndufs4*^*-/-*^ mice decreases from 64 days to 29 days in *Ndufs4*^*-/-*^ *Nes-Phd2*^*-/-*^ triple transgenic mice that display Hif activation (*34*). Increased iron absorption via hyperactivation of hypoxia inducible factors may partially explain this surprising result.

## Conclusion

This work establishes that mice missing the iron-sulfur cluster containing ETC subunit NDUFS4 have iron overload in the liver, and that iron restriction is sufficient to delay the onset of disease progression. Proteins sensitive to changes in intracellular iron, such as TFR1 and ferritin, change in response to this iron overload. This is consistent with an increase in labile iron that can be recognized by iron response proteins and is commonly associated with iron-dependent oxidative stress. Iron restriction causes anemia, decreases liver iron levels, reduces MDA formation, and subsequently alters the expression of these iron regulatory proteins.

Iron deposition in multiple tissues, including the brain, with aging is thought to contribute to the development of age-related diseases (*36, 49*). Iron dyshomeostasis in the brain is an early feature of cognitive decline, such as in Parkinson’s Disease, and contributes to neuroinflammation, proteostasis failure, and other physiological abnormalities. There remains an unmet need to deeply understand the influence of mitochondrial dysfunction on iron homeostasis in normative aging and neurodegeneration. Our data suggest that decreased Complex I activity with aging may help contribute to this neurodegeneration with brain iron accumulation, and therapeutic interventions targeting genetic or age-progressive causes of mitochondrial dysfunction may be especially beneficial.

The mitochondrial disease Leigh Syndrome is caused by mutations in more than 75 nuclear- or mitochondrial-encoded genes such as several subunits of Complex I (e.g. NDUFS4), Complex IV (e.g. COX10), assembly factors (e.g. SURF1), metabolic enzymes (PDHA1), and transport proteins (SLC19A3) amongst others (*1*). It is critical to better understand the generality of pathogenic iron dyshomeostasis in Leigh Syndrome patients with these specific mutations. This work may have broader implications in the development of diagnostic markers for Leigh Syndrome, in which serum iron, transferrin saturation, serum ferritin, or serum hepcidin can be readily analyzed clinically. It may also lead to further beneficial treatment regimens in Leigh Syndrome or other mitochondrial diseases.

## Materials and Methods

### Key Resources Table

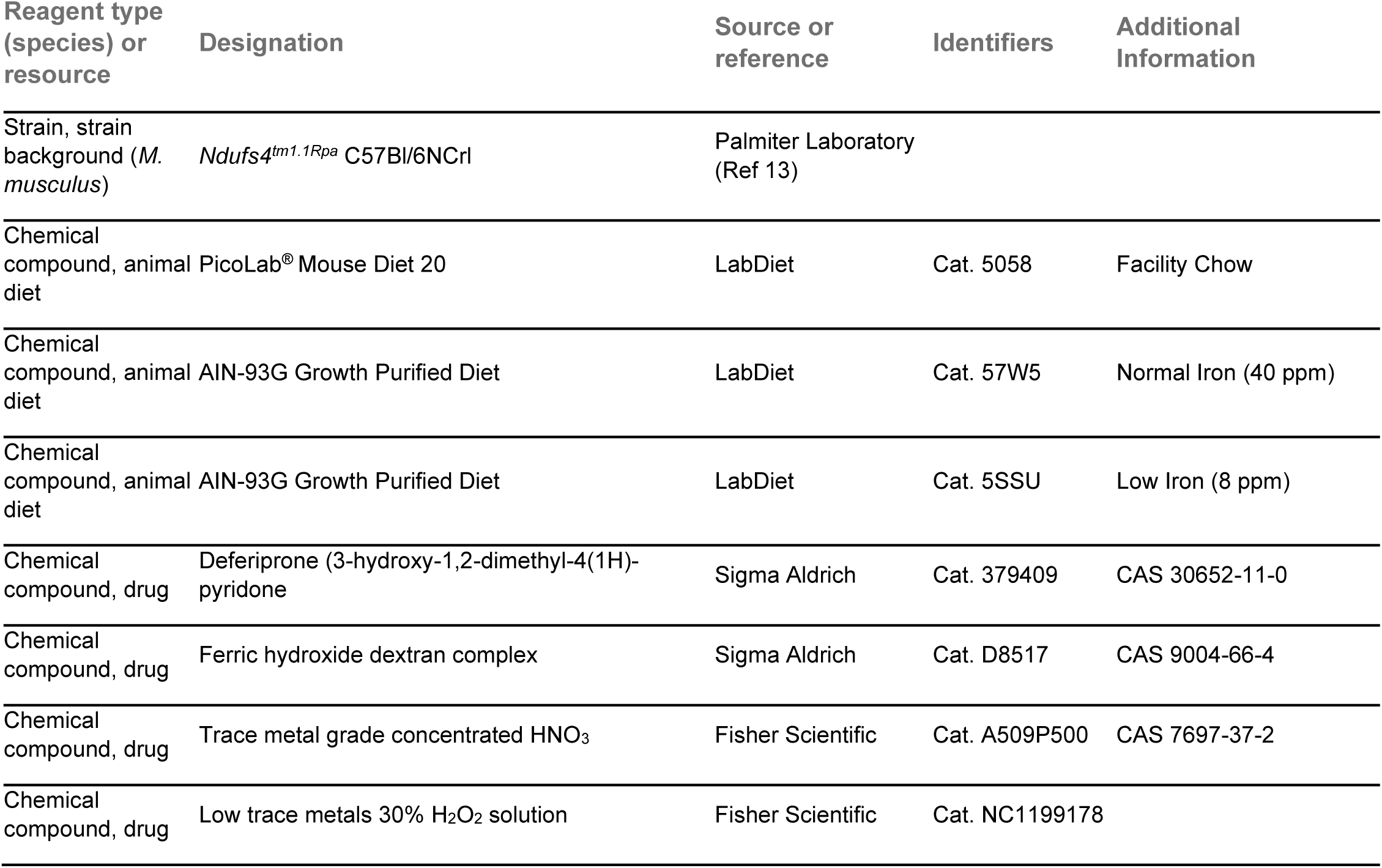
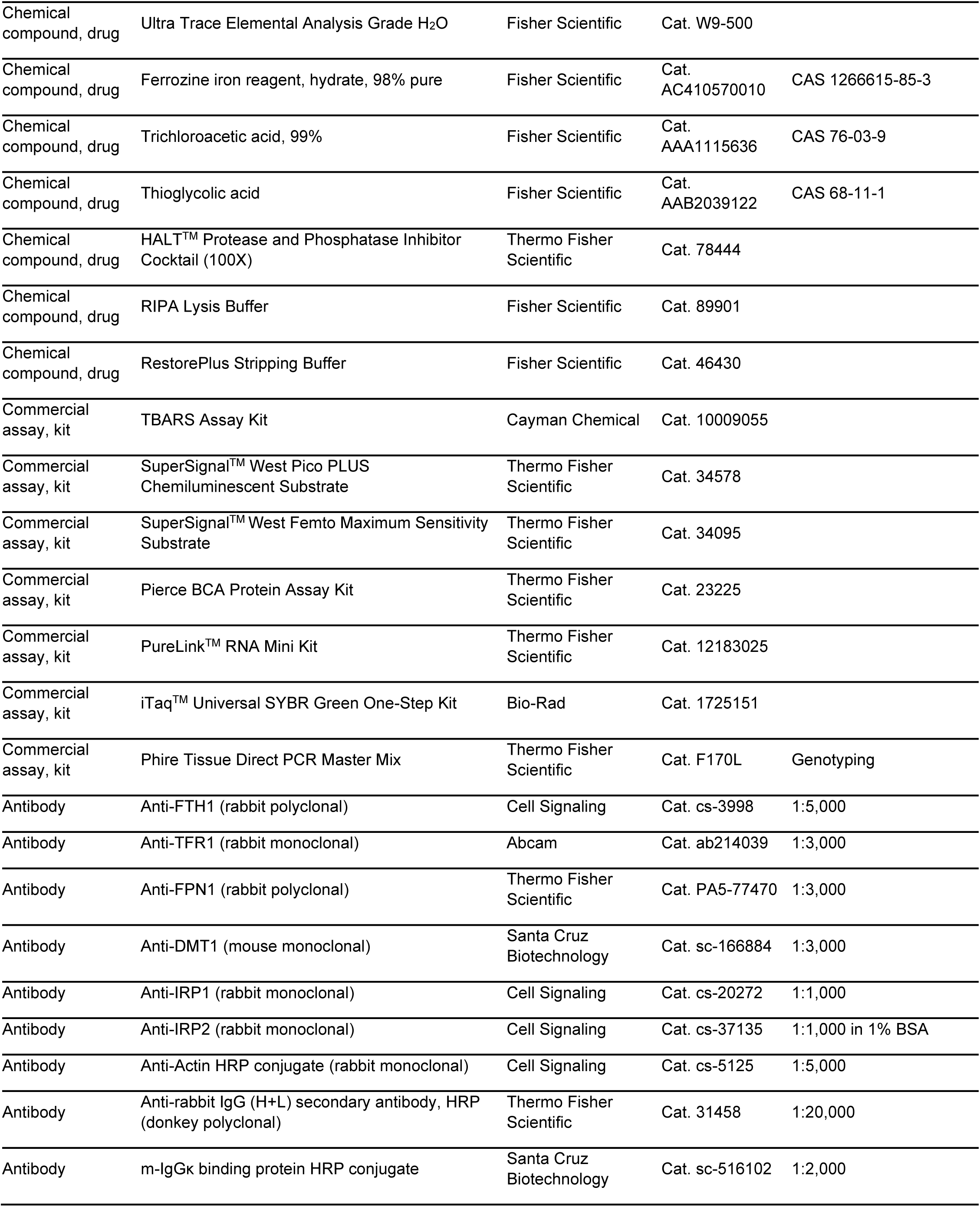
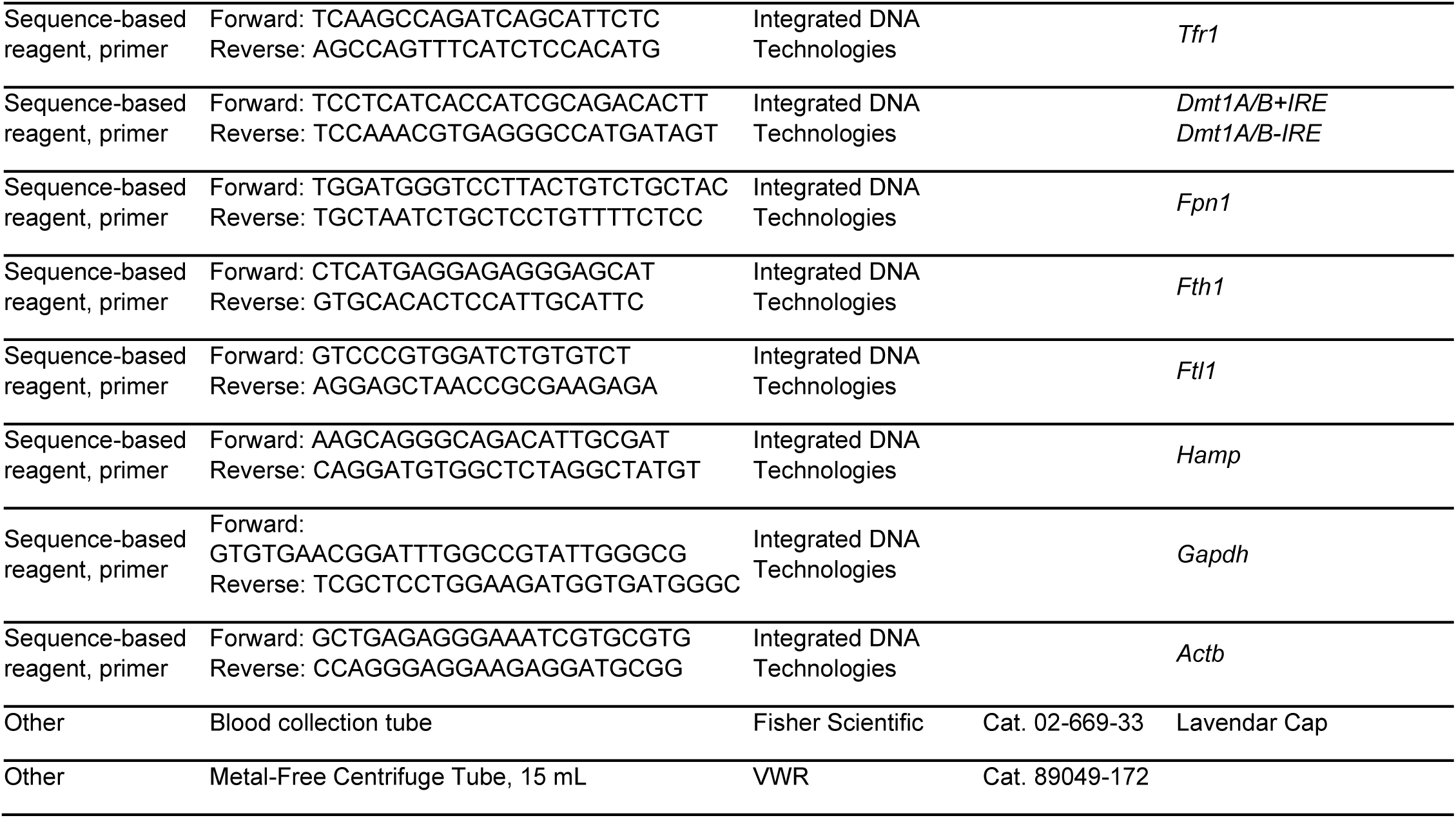

### Animals and Animal Use

Experiments, procedures, and protocols described herein to care for, and handle, mice were reviewed and approved by the University of Washington Institutional Animal Care and Use Committee (IACUC) and strictly adhered to guidelines described in the Guide for the Care and Use of Laboratory Animals of the National Institutes of Health. Mice were housed at the University of Washington on a 14:10-hour light/dark cycle and provided food, water, and/or gel *ad libitum. Ndufs4*^*+/-*^ mice were donated by the Palmiter Laboratory at the University of Washington and serially backcrossed with C57Bl/6NCrl mice. Pups produced by pairwise breeding of *Ndufs4*^*+/-*^ mice were genotyped by PCR before post-natal day 21 (PND21) to identify *Ndufs4*^*+/+*^, *Ndufs4*^*+/-*^, and *Ndufs4*^*-/-*^ pups. Male and female mice were distributed evenly in experiments, and *Ndufs4*^*-/-*^ mice were housed with at least one healthy littermate for companionship and warmth. No more than 5 mice were housed per cage. Mice were monitored daily for common signs of neurodegeneration (e.g. clasping), weight loss, and moribundity. Mice that reached endpoint criteria, including >30% decrease from maximum body weight, loss of righting reflex, or were unresponsive to stimuli, were euthanized via primary and secondary CO_2_ asphyxiation.

Mice were weaned around P21 and fed standard facility chow (PicoLab^®^ Mouse Diet 20 5058) or synthetic AIN-93G Growth Purified Diet containing normal (40 ppm, Lab Diet 57W5) or low (8 ppm, LabDiet 5SSU) amounts of iron. Mice fed the chow diet were treated with deferiprone in the water (0.5, 1, or 2 mg/mL water). Food and water were replenished every ∼5 days. Mice on the control synthetic AIN-93G diet were injected every 3 days via intraperitoneal (i.p.) injection with iron-dextran (100 mg/kg body weight in saline) that was made from a stock containing 100 mg/mL iron. The volume injected was 6.67 µL/g body weight and mice were injected on alternate sides to minimize irritation or distress.

### Statistical Analysis

The data presented in this study represents the weighted mean ± SEM with a minimum of three biological replicates and multiple technical replicates whenever possible. All studies utilized both male and female mice. Statistical analysis for lifespan data was performed using a log-rank test, and P-values for other experiments were calculated using student t-test with Bonferroni correction or with an ANOVA test where appropriate. Pearson’s test was used to test significance of correlations. * p<0.05, ** p<0.01, *** p<0.001, **** p<0.0001

### Tissue Collection

Tissues were freshly collected for protein, mRNA, and metabolic experiments from non-fasted ∼35-day old mice in the morning. Mice were euthanized via cervical dislocation and tissues were excised and flash frozen in liquid N_2_ before processing. Tissues for analysis of metal content were obtained and flash frozen from non-fasted ∼35-day old mice after anesthetization with a ketamine/xylazine cocktail and blood exhaustively removed via whole body perfusion with PBS.

### Complete Blood Count

Blood was collected via cardiac puncture from non-fasted ∼30-day old mice after anesthetization with a ketamine/xylazine cocktail. Blood was immediately transferred to a lavender cap blood collection tube containing EDTA to prevent coagulation. The blood was stored on ice and immediately submitted to the Veterinary Diagnostics Laboratory at the University of Washington for CBC analysis.

### Analysis of Total Metal Content via ICP-MS

Tissues were collected after extensive perfusion as described above in non-fasted PND35 mice and flash frozen in liquid N_2_. Tissues were thoroughly homogenized in liquid N_2_ and aliquoted into pre-tared metal free Eppendorf tubes. Tissues were dried via a SpeedVac vacuum concentrator and weighed. Tissues were then digested with HNO_3_/H_2_O_2_ after slowly heating to 95 °C overnight. Digestates were then diluted in trace metal free water containing dilute HNO_3_ and submitted to the trace element analysis laboratory at the University of Washington for ICP-MS quantification. Any glassware or non-trace metal grade containers were washed with HCl acid solution to remove trace metals.

### Quantification of Non-Heme Iron with Ferrozine

A modified ferrozine-based assay (*52*) was used to quantify levels of non-heme iron in PND35 livers. Either standards (prepared from FeSO_4_) or homogenized liver tissue were digested with 3M HCl containing 600 mM trichloroacetic acid (TCA). Samples were heated at 65 °C for 20 hours, centrifuged, and supernatant collected. Chromogen solution (100 µL containing 500 µM ferrozine, 1.5M sodium acetate, 1.5% v/v thioglycolic acid in trace metal free H_2_O) was added to digestate (50 µL) in a 96-well plate and incubated for 30 minutes at room temperature to form a purple-colored solution. Ferrozine was quantified via measuring absorbance at 595 nM with a microplate reader and non-heme iron was calculated using the obtained standard curve.

### Quantification of MDA-TBA Adduct

Lipid peroxidation was assessed by a malondialdehyde thiobarbituric acid (MDA-TBA) reactive substance assay using a modified procedure from the Cayman Chemicals MDA-TBA Assay Kit. Homogenized livers from PND35 mice were lysed using RIPA buffer containing protease inhibitors and supernatant collected after centrifugation. MDA-TBA adduct was prepared according to manufacturer instructions, except we used half the recommended volumes to increase signal. We read the absorbance of the MDA-TBA adduct at 530 nM and calculated MDA-TBA concentrations using a standard curve.

### Protein Analysis via Western Blot

Proteins were extracted from homogenized liver and brains from PND35 mice with RIPA buffer containing protease and phosphatase inhibitors. Protein extract was collected after centrifugation, quantified by a BCA assay, and diluted to either 1 or 2 mg/mL. Samples for western blot were prepared by diluting protein (20 µg) in Laemmli sample buffer, reducing agent, and RIPA buffer. Samples were denatured at 95 °C for 5 minutes and loaded onto NuPAGE 4-12% MIDI or MINI gels and ran at 120 V in MOPS running buffer. Protein was transferred to a PVDF membrane using a Trans-Blot^®^ Turbo Transfer system with manufacturer transfer buffer containing an additional 0.2% SDS. Blots were blocked with either 5% BSA or 5% milk in TBST at room temperature for 1 hour. Membranes were incubated in either 1% or 5% BSA with primary antibodies overnight at 4 °C, followed by 2 hours at room temperature. Membranes were rinsed with TBST, and incubated with secondary antibody for 1-2 hours in 5% BSA or 5% milk. After rinsing membranes with TBST, either Thermo Fisher SuperSignal PicoPlus or Femto ECL reagent was added and chemiluminescence detected with an iBright CL1500 system. Membranes were stripped for 10 minutes, rinsed, blocked, and re-probed as described above. Densitometry was performed using the iBright Analysis Software relative to actin loading control, and normalized to wild type levels.

### mRNA Quantification via qRT-PCR

Total RNA was isolated from PND35 livers using the Thermo Fisher PureLink™ RNA Mini Kit according to manufacturer instructions and quantified with a NanoDrop spectrophotometer. Primers reported from the literature were checked by NCBI Primer Blast for accuracy. Primers were purchased from Integrative DNA Technologies as RxnReady Oligos and diluted to a 1 mM stock. Expression of mRNA was quantified by qRT-PCR using the BioRad iTaq™ Universal SYBR^®^ Green One-Step RT-PCR Kit containing 300 nM primer and 50 ng/µL of total RNA in a 10 µL reaction. PCR reaction products were verified by melting temperature or gel electrophoresis and compared to expected band size. mRNA expression relative to *Gapdh* or *Actb* was normalized to wild type levels.

## Acknowledgements

We thank Sarah Proffitt and Kerrie Allen for assistance with CBC analysis through the University of Washington Veterinary Diagnostics Laboratory. We thank Alex Gagnon and Tamas Ugrai from the University of Washington Trace Element Laboratory for Environmental Science for quantification of metal content by ICP-MS. We gratefully acknowledge Will Rieger, Tom Milstein, and Sanchita Narayan for assistance with animal husbandry.

## Author Contributions

ASG and MK conceptualized and designed the study. ASG, CK, VTH, CMB, RKC, NTH, SH, HDK, AOM, JS, and Y-CP performed the experiments. ASG, CK, VTH, and CB analyzed the data. ASG wrote the manuscript. ASG, CK, and MK edited the manuscript.

## SUPPLEMENTAL DATA AND FIGURES

**Figure supplement 1.**
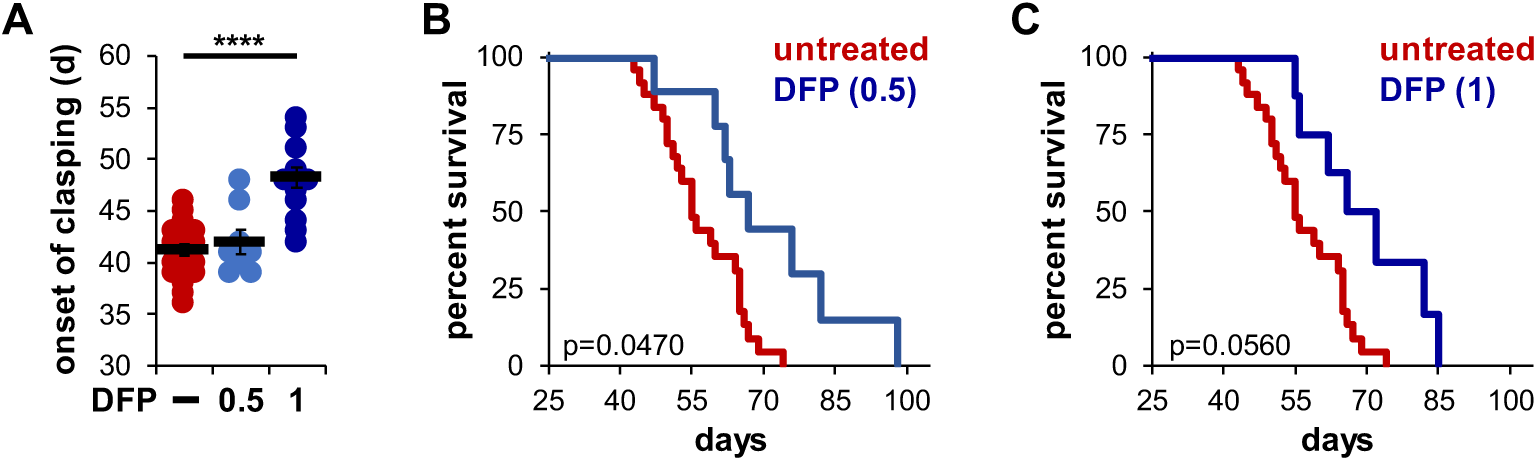
(A) Age at which *Ndufs4*^*-/-*^ mice exhibited the clasping phenotype on chow diet. Mice were treated with either vehicle or deferiprone (DFP) in the water (0.5 or 1 mg/mL) from weaning. (B) Survival curves of *Ndufs4*^*-/-*^ mice fed a chow diet and treated with deferiprone in the water (0.5 mg/mL) from weaning. (C) Survival curves of *Ndufs4*^*-/-*^ mice fed a chow diet and treated with deferiprone in the water (1 mg/mL) from weaning. p-value was calculated by log-rank for lifespan analyses. **** p<0.0001, t-test.

**Figure supplement 2.**
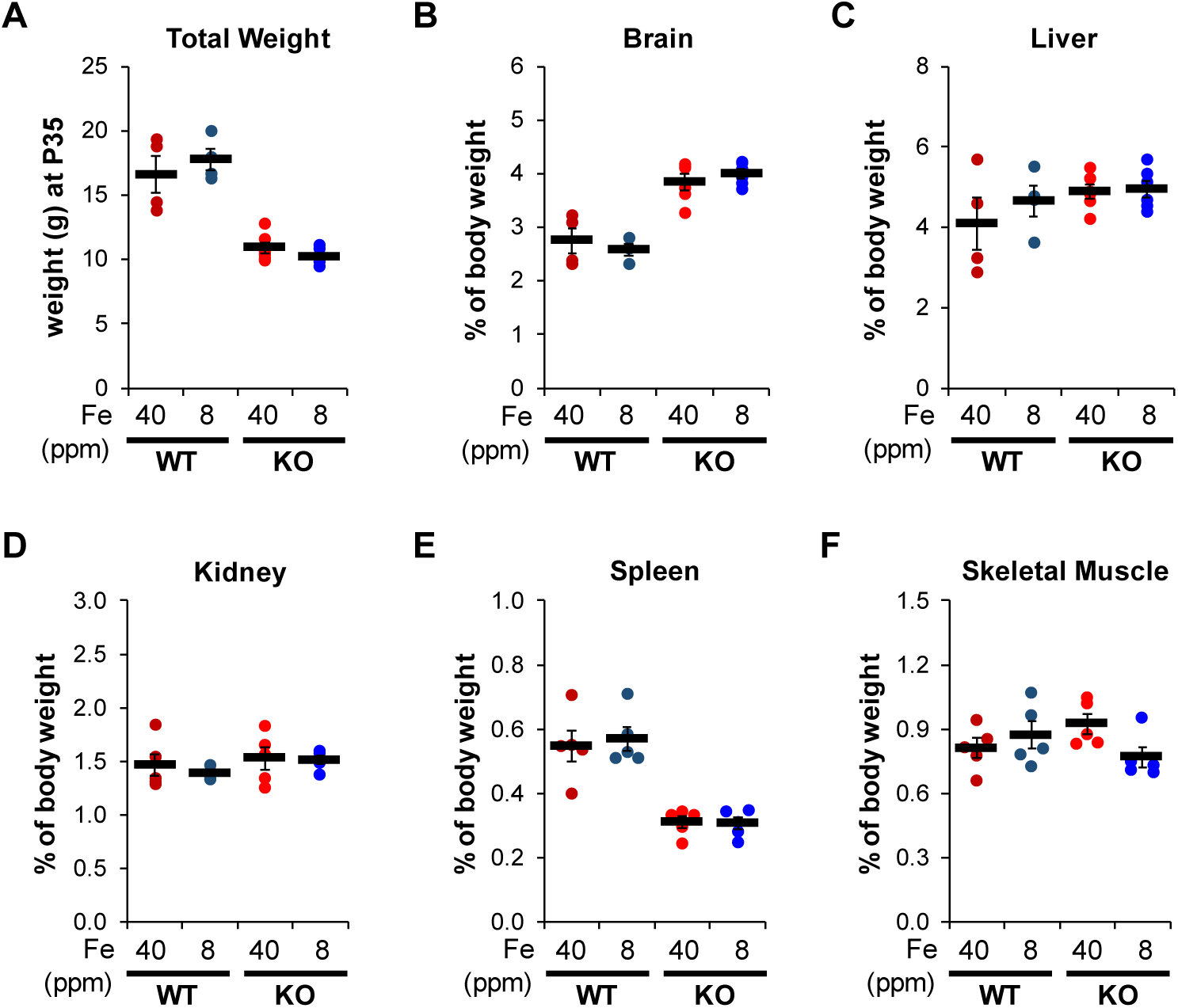
(A) Weight at time of tissue collection (PND35) of WT or *Ndufs4*^*-/-*^ mice on a normal (40 ppm) or low (8 ppm) AIN-93G synthetic diet. (B) Percent of tissue weight from mice in (A) relative to total body weight at time of collection (PND35) in brain, (C) liver, (D) kidney, (E) spleen, and (F) quadricep.

**Figure Supplement 3.**
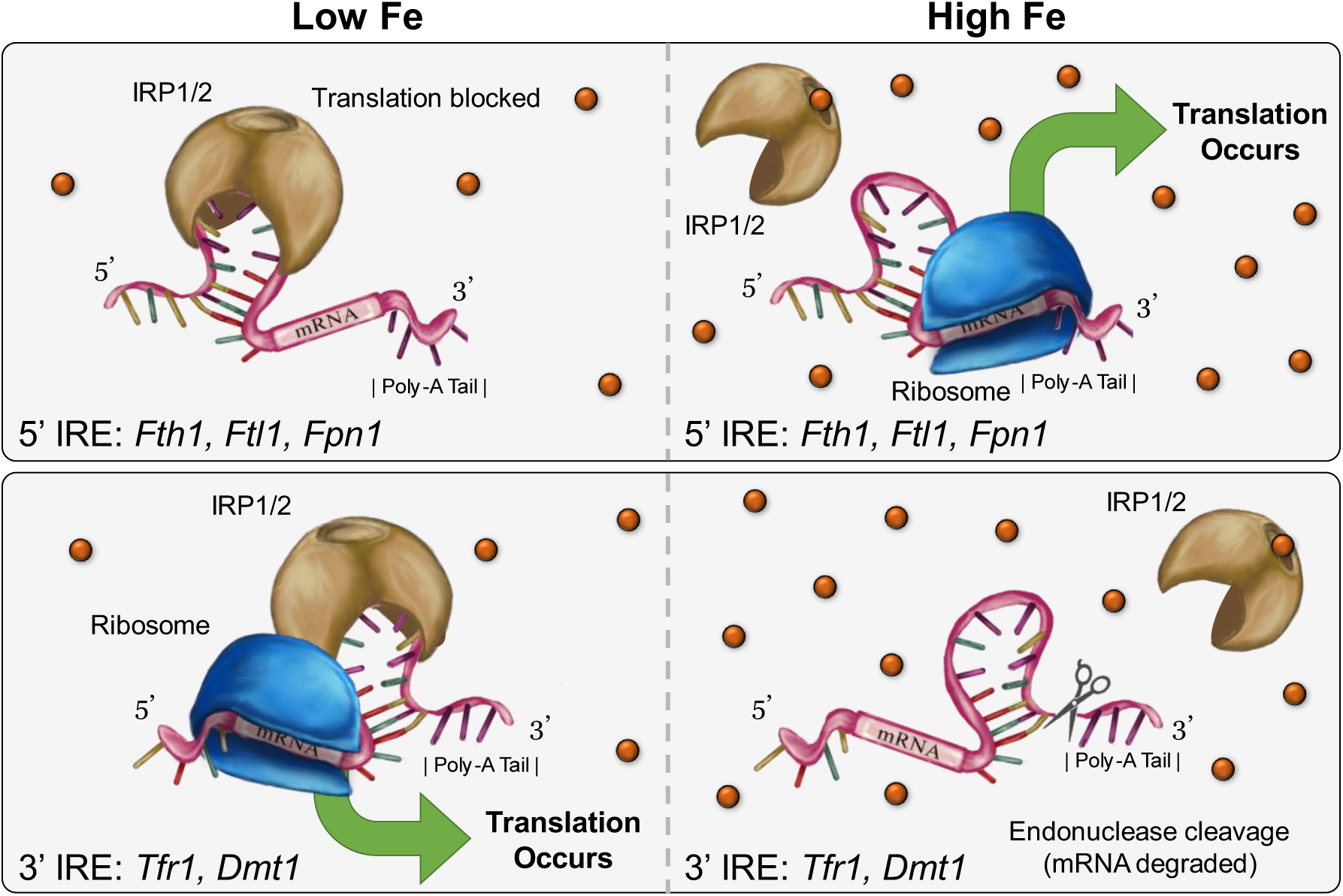
Simplified schematic of IRE-dependent translational regulation of proteins involved in iron transport, storage, or metabolism. Translation is blocked for 5’-IREs (e.g. *Fth1, Ftl1, Fpn1*) when iron regulatory proteins are bound to the IRE in low iron conditions (top left). Protein expression increases due to mRNA stabilization when iron regulatory proteins are bound to 3’-IREs (e.g. *Tfr1, Dmt1*) in low iron conditions (bottom left). Exposure to high iron allows for translation in genes containing 5’-IREs (top right) and leads to decreased protein expression for 3’-IREs due to endonuclease-mediated mRNA degradation (bottom right).

**Figure supplement 4.**
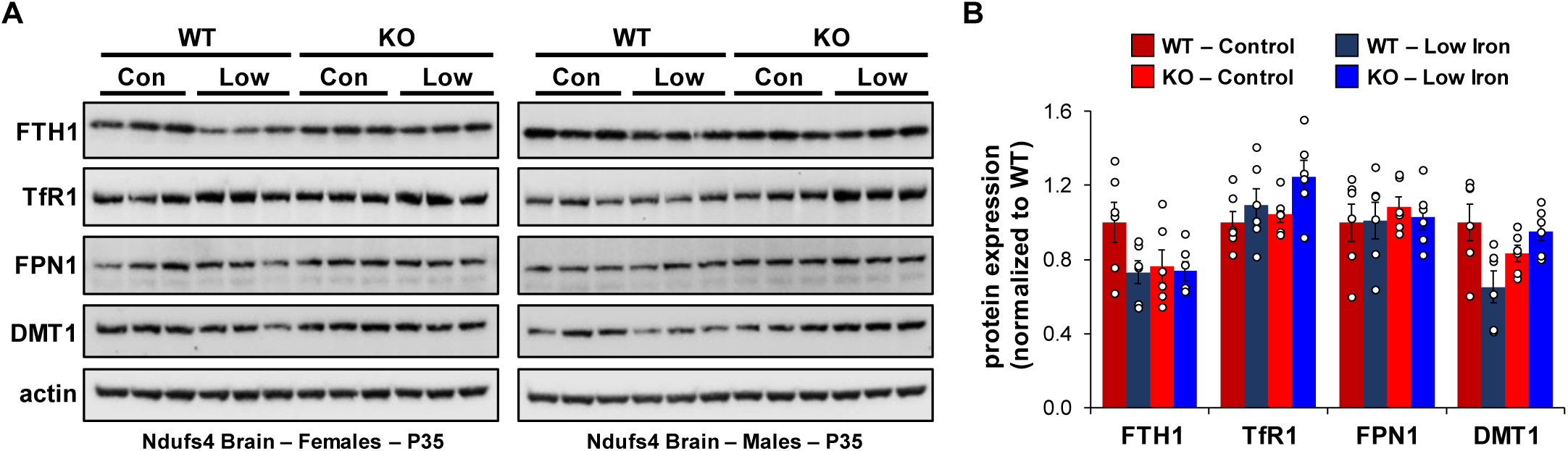
(A) Representative western blot images and (B) densitometry (relative to actin) of proteins involved in regulation of iron transport, storage, or metabolism in whole brains from PND35 WT and *Ndufs4*^*-/-*^ mice fed a control (40 ppm) or low (8 ppm) iron AIN-93G synthetic diet from weaning. Each lane represents protein extract from a single mouse.

